# Stimulus induced visual cortical networks are recapitulated by spontaneous local and inter-areal synchronization

**DOI:** 10.1101/021055

**Authors:** C. M. Lewis, C. A. Bosman, T. Womelsdorf, P. Fries

## Abstract

Intrinsic covariation of brain activity has been studied across many levels of brain organization. Between visual areas, neuronal activity covaries primarily among portions with similar retinotopic selectivity. We hypothesized that spontaneous inter-areal co-activation is subserved by neuronal synchronization. We performed simultaneous high-density electrocorticographic recordings across several visual areas in awake monkeys to investigate spatial patterns of local and inter-areal synchronization. We show that stimulation-induced patterns of inter-areal co-activation were reactivated in the absence of stimulation. Reactivation occurred through both, inter-areal co-fluctuation of local activity and inter-areal phase synchronization. Furthermore, the trial-by-trial covariance of the induced responses recapitulated the pattern of inter-areal coupling observed during stimulation, i.e. the signal correlation. Reactivation-related synchronization showed distinct peaks in the theta, alpha and gamma frequency bands. During passive states, this rhythmic reactivation was augmented by specific patterns of arrhythmic correspondence. These results suggest that networks of intrinsic covariation observed at multiple levels and with several recording techniques are related to synchronization and that behavioral state may affect the structure of intrinsic dynamics.

## Introduction

A classical approach to brain research considers the brain as responding to external stimuli and converting them into appropriate behavioral output. In this framework, endogenously driven variance in the brain is considered noise (Shadlen and Newsome, 1994). However, this so-called noise has been found to be highly structured and influenced by behavioral context (Cohen and Newsome, 2008). Such variation is a flavor of the ongoing, spontaneous activity of the brain. Both, the variation in the brain’s response to identical stimulation and spontaneous activity in the absence of stimulation are endogenously generated and structured in spatially specific intrinsic networks. Highly specific intrinsic networks have been described at essentially all spatial scales: from two individual neurons up to the whole brain. Membrane potentials of nearby neurons show a high degree of spontaneous correlation (Lampl et al., 1999). Neuronal spike rates co-fluctuate across physically identical trials, and this so-called noise-correlation between neurons is related to the similarity in their stimulus selectivity (Luczak et al., 2009). Correspondingly, when entire maps of population activity are investigated, patterns of activation induced by stimuli are found to re-occur spontaneously (Kenet et al., 2003). The same holds across neighboring maps in auditory cortex, where population activity spontaneously reproduces the tonotopic organization (Fukushima et al., 2012). Such co-activations can also be observed with fMRI. Across visual areas, regions selective for either foveal or peripheral stimuli show correlated BOLD activity (Vincent et al., 2007). In human subjects, BOLD signals were found to covary in symmetric, bilateral foci in the two hemispheres (Biswal et al., 1995). Subsequently, brain-wide sets of areas that were typically co-activated or co-deactivated by specific cognitive tasks were described (Fox and Raichle, 2007; Greicius et al., 2003). The finding of robust intrinsic networks using fMRI led to a large number of studies (Fox and Raichle, 2007), which have recently demonstrated the functional importance of intrinsic networks. For example, spontaneous, correlated BOLD signals predict behavior (Hesselmann et al., 2008), and affect learning (Baldassarre et al., 2012; Lewis et al., 2009).

Most fMRI studies of intrinsic networks involve inter-areal correlations, yet we have only a partial understanding of the underlying mechanisms. Recently, it has been shown that inter-areal BOLD signal correlations between regions of Squirrel monkey somatosensory cortex reflect the somatotopic map and are subserved by millisecond scale spike correlations (Wang et al., 2013). The rhythmic synchronization of neuronal activity is an interesting candidate mechanism for inter-areal interactions. Local rhythmic synchronization likely enhances neuronal impact through coincident postsynaptic input (Fries et al., 2001; Salinas and Sejnowski, 2001). Furthermore, inter-areal rhythmic synchronization aligns temporal windows of excitability and likely renders communication effective (Fries, 2005; Womelsdorf et al., 2007).

We investigate here whether rhythmic synchronization subserves intrinsic networks between visual areas. Using high-resolution multi-area electrocorticography (ECoG) in awake monkeys, and utilizing the retinotopy of early and intermediate visual areas, we show that intrinsic inter-areal networks recapitulate stimulus induced inter-areal networks, and are subserved by local and inter-areal synchronization in the theta, alpha and gamma-frequency bands. Surprisingly, even though no clear peaks were evident in power spectra of spontaneous activity, intrinsic networks found in the absence of visual stimulation revealed both arrhythmic and rhythmic processes. Therefore, specific patterns of rhythmic synchronization can exist in power spectra without clear peaks (Vinck et al., 2014). Further, the emergence of arrhythmic retinotopic coactivation, specifically in passive behavioral states may indicate a distinct dynamical mode. The results suggest that transitions between behavioral state shape intrinsic networks. Overall, our findings provide a conceptual bridge by linking intrinsic networks to the insights that have been obtained about the biophysical mechanisms underlying rhythmic and arrhythmic synchronization.

## Results

We investigated whether intrinsic networks between early (areas V1 and V2) and intermediate (areas V4 and TEO) visual regions are subserved by rhythmic activity and synchronization. To this end, we obtained measurements of neuronal activity with millisecond temporal resolution and few-millimeter spatial resolution and simultaneous coverage of both cortical regions. We employed electrocorticography (ECoG) in two awake macaque monkeys (Monkey P, Fig 1A; Monkey K, Fig S1A). For all analyses shown here, the signals from immediately neighboring ECoG electrodes were subtracted from each other to obtain local bipolar derivatives, which are free of the common recording reference, and to which we will refer to as “sites”. We explored intrinsic networks arising from covariations in local synchrony, i.e. correlated variation of endogenous power between areas, or in inter-areal synchronization, i.e. coherence between areas. As intrinsic networks can emerge spontaneously, in the absence of stimulation, or as trial-by-trial variation in stimulus-induced responses, we examined the correspondence of the spatial structure of the observed networks to the known spatial structure of visual cortex, namely retinotopy. If the observed intrinsic power covariation or coherence is spatially correlated to retinotopy, it is unlikely to be noise and may have a functional role.

**Figure 1.**
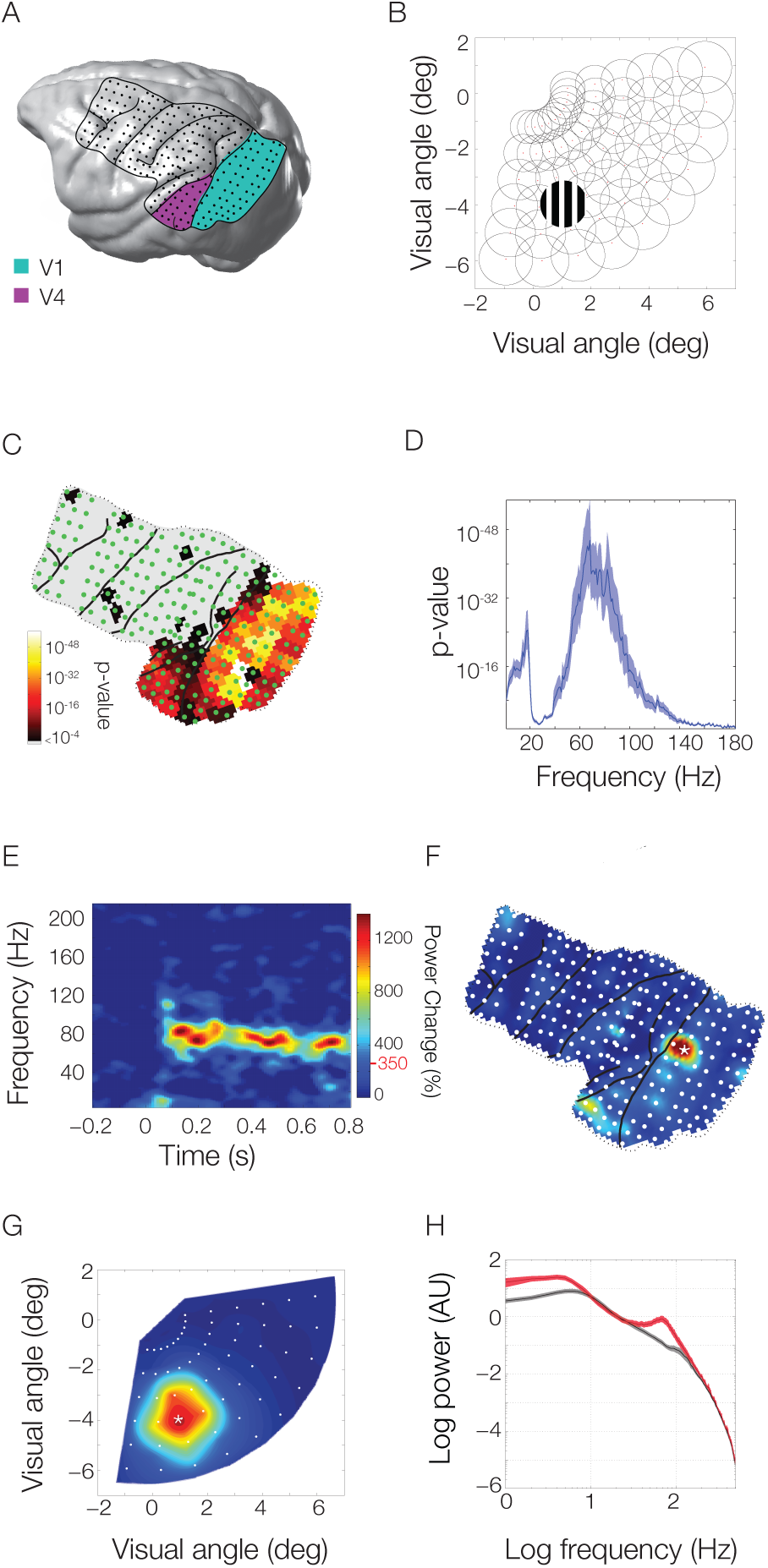
High density ECoG layout and receptive field mapping paradigm. (A) Rendering of the brain of monkey P with the ECoG grid overlaid. Lines indicate the covered area with the major sulci. Dots indicate the 218 bipolar electrode derivations. Sites considered as lying in areas V1 and V2 are highlighted in green and those in areas V4 and TEO are highlighted in purple. (B) Receptive fields were mapped with scrolling gratings at 60 locations in the lower right quadrant, corresponding to the coverage of the ECoG array. (C) Selectivity of all ECoG sites for stimulus position based on stimulus induced power in all frequency bands. (D) Selectivity of position tuned sites as a function of frequency. (E-G) Example average response to stimulation at the position marked in (B). (E) Time-Frequency plot at the site marked by a star in (F). Topographic plot of induced gamma power (80-95 Hz) for each ECoG site. (G) Induced gamma band response across all positions for the site marked in (F). Colorbar is the same for (E-G), red line and value next to colorbar indicate significance.

## Retinotopic selectivity of ECoG signals

To assess retinotopic selectivity across all sites of the ECoG grid, monkeys kept fixation for several seconds while visual stimuli were presented randomly interleaved at 60 different positions in the lower right visual quadrant (Fig 1B), corresponding to the portion of visual space covered by our grid. Stimulus-position dependent changes in spectral power (in any frequency band) were established through an Analysis of Variance (ANOVA). The resulting *p*-values are shown in figure 1C (for monkey P; for monkey K, see Fig. S1B) and reveal that retinotopy was mainly found in areas V1/V2 and areas V4/TEO. We selected those sites for further analyses (V1/V2: 68 sites in monkey P, 32 in monkey K; V4/TEO: 17 sites in both monkeys). For these sites, figure 1D shows how well stimulus-position was distinguished based on stimulus-induced power as a function of frequency (both monkeys combined, individual spectra are shown in Fig. S2A and B). This demonstrates that retinotopic selectivity was present predominantly in a low frequency band (1-20 Hz) and a gamma-frequency band (60-100 Hz). These two bands also contained the largest stimulus induced power, as is illustrated in the time-frequency analysis of one example site from area V1 activated by its optimal stimulus (Fig. 1E monkey P, for monkey K see Fig. S1C). This pattern of activity was remarkably consistent across recording sites, stimulus positions and monkeys (Fig. S2). While low frequency activity was present both with and without stimulation, gamma-band activity was primarily stimulus induced and had the highest degree of retinotopic selectivity. Correspondingly, we illustrate the gamma-band power for all ECoG sites for the same stimulus position in Fig. 1F (monkey P, for monkey K see Fig. S1D). The well-localized topographical activation suggests that a given site responds only to a select region of visual space, as was previously shown for these data (Bosman et al., 2012). Indeed, when the site with maximal response in figure 1F (marked with a star) was selected and the response to all stimulus locations was displayed in figure 1G (monkey P, for monkey K, see Fig. S1E), there was a clearly defined receptive field (RF).

To further investigate the retinotopy of the ECoG recordings from V1, V2, V4 and TEO, we grouped the 60 stimulus locations according to eccentricity or elevation. Both eccentricity (Fig. 2A for monkey P; for monkey K, see Fig. S3A) and elevation (Fig. 2B for monkey P; for monkey K, see Fig. S3B) were represented in orderly retinotopic maps, that corresponded well with previously determined topographies from repeated recordings with penetrating electrodes (Gattass et al., 2005) or from fMRI (Brewer et al., 2002). Two contiguous maps of space were visible, one behind the lunate sulcus for areas V1/V2, and another one between the lunate and the superior temporal sulcus for areas V4/TEO. For simplicity, we will refer to ECoG sites in the V1/V2 map as V1, and to sites in the V4/TEO map as V4. We investigated whether these maps determined the intrinsic power covariations and/or the coherence between areas. A schematic of our processing stream is displayed in figure 3A. The main steps are: 1) Quantify the spatial pattern of stimulus induced power correlation (the covariation of V1–V4 sites by stimulus position); 2) Quantify the spatial pattern of intrinsic power correlation or coherence; 3) Correlate the spatial patterns from 1) and 2). As both stimulus induced (1) and intrinsic (2) metrics could be estimated for a full spectrum of frequencies, the correlation (3) could be determined for all pairs of frequencies. In order to investigate the full pattern of reactivation, we quantified the correspondence between intrinsic synchronization and retinotopy across all frequency pairs. We could therefore document the rich pattern of inter-areal synchronization in a complete and unbiased manner.

**Figure 2.**
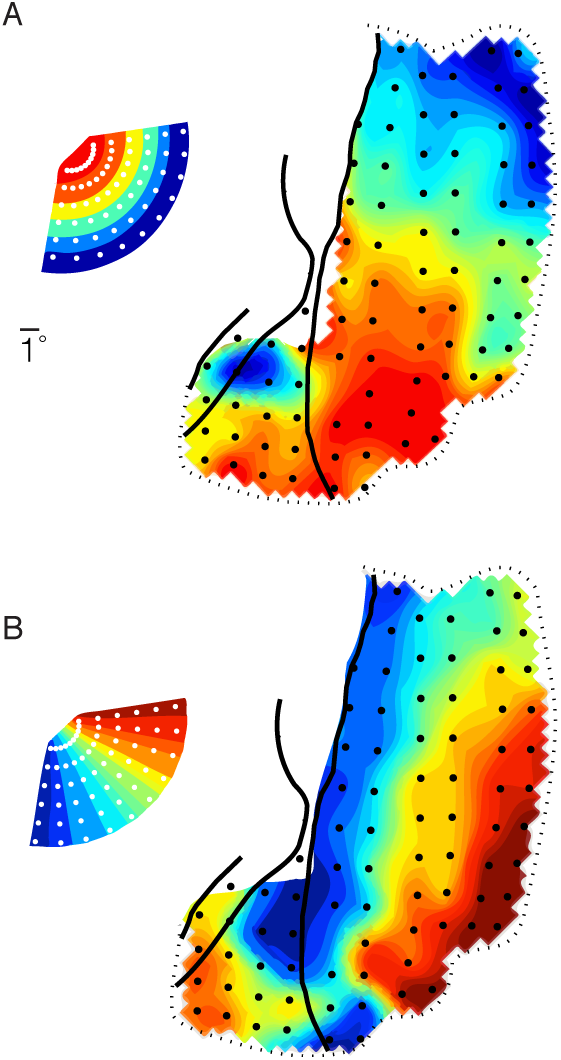
Retinotopic maps based on gamma band (80-95 hz) activity in monkey P. (A) Map of eccentricity; each recording site is colored to indicate the mean eccentricity of the 5 stimuli giving the largest gamma band response. (B) Map of elevation; each recording site is colored to indicate the mean elevation as estimated above. Inset shows how the 60 stimulus locations are represented across eccentricity and elevation.

**Figure 3.**
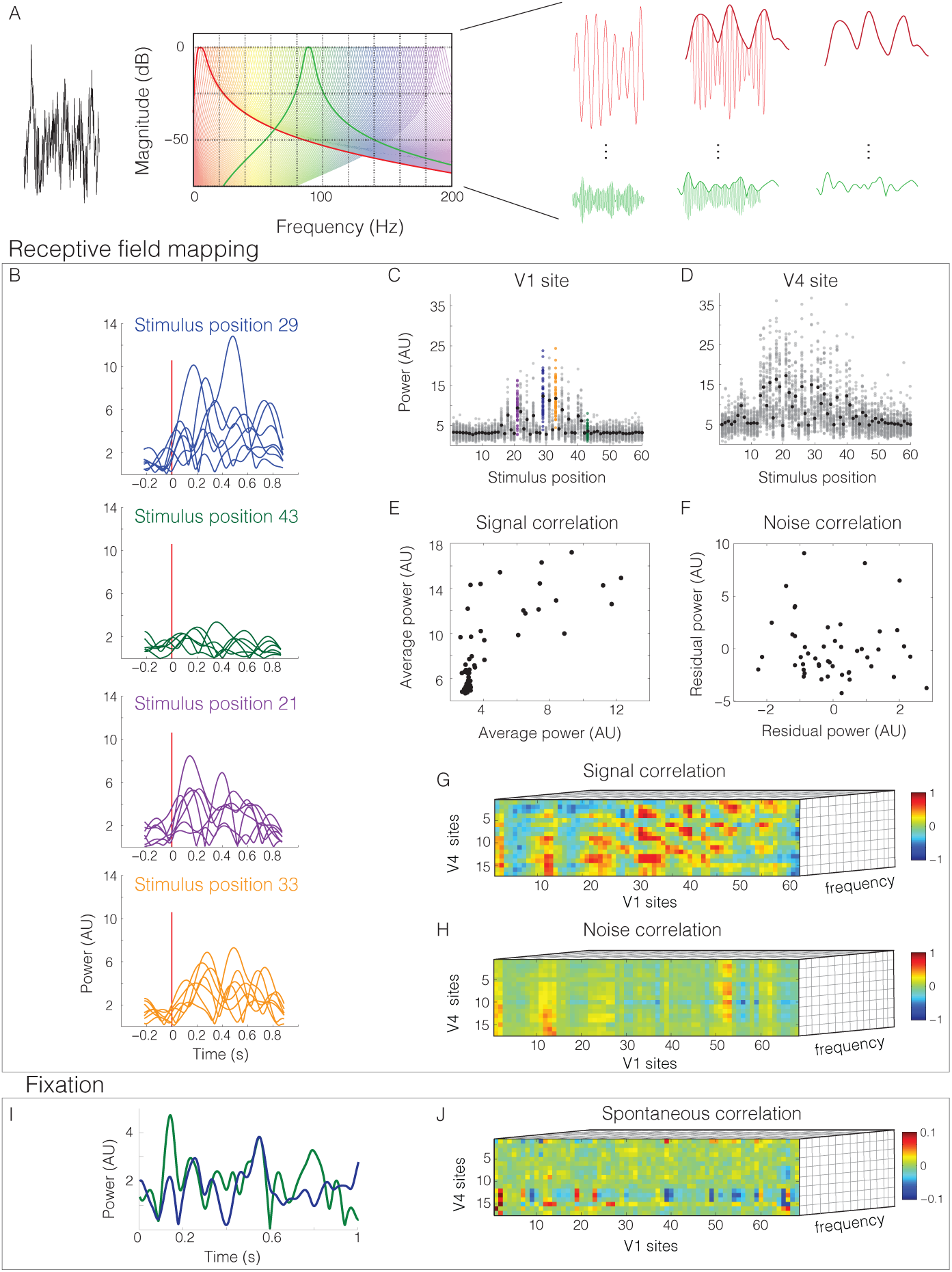
Analysis pipeline for visual stimulation and passive fixation. (A) Signal processing pipeline: Raw field potentials (left) were filtered in 185 linearly spaced bands between 2 Hz and 200 Hz (center). The power envelope was extracted from the analytic Hilbert representation (right). Receptive field mapping. (B) Example power responses in the gamma band (80-95 Hz) for one V1 site to 7 repetitions of visual stimulation at 4 positions. (C) Average (in black) and individual trial (in grey) responses of the same V1 site to all 60 stimulus positions. Positions in (B) shown in the respective color. (D) Same as (C) but for an example V4 site. (E) The signal correlation for the two sites shown above. (F) The noise correlation for the same two sites. Signal (G) and Noise (H) correlations were computed between all pairs of V1-V4 sites for all frequencies of interest. Passive fixation. (I) One second of activity in the gamma band for the same V1-V4 sites shown above. (J) Spontaneous correlation was computed for all pairs of V1-V4 sites in each frequency of interest.

## Signal Correlations

In order to assess the similarity of stimulus preferences between V1 sites and V4 sites, we computed for each inter-areal site pair the correlation between the average induced power per stimulus position, across the different stimulus positions. Figure 3B shows the gamma band (80-95 Hz) response of a single V1 site to 7 repetitions of visual stimulation in 4 different stimulus positions. The gamma band response varied systematically as a function of stimulus position in both V1 (Fig. 3C) and V4 (Fig. 3D), as expected from the results shown in Figures 1 and 2. It is common (Gawne and Richmond, 1993) to decompose the response to a stimulus into a signal component, considered to be the average response to multiple identical stimulus presentations (black dots in Fig. 3C for a V1 site and D for a V4 site), and a noise component, considered to be the difference between the average response and the actual response to a particular stimulus presentation (variation of the grey dots in Fig. 3C and D around the black dots). In this framework, the correlation between two sites’ average stimulus-induced power across different stimuli is considered the signal correlation (sample site pair shown in Fig. 3E), and we will use this term in the following. The signal correlation is calculated per inter-areal site pair and per frequency band, across the 60 stimulus positions (Fig. 3G). The signal correlation captures the similarity of the two sites’ stimulus selectivity and therefore reflects the underlying retinotopy. Figure 4A shows the matrix of signal correlations for all possible V1-V4 site pairs for the gamma-frequency band (between 80 and 95 Hz) for monkey P. The repetitive structure in the matrix along both axes reflects the arrangement of electrodes on both areas in distinct lanes, which have been unwrapped along the x- and y-axes. This matrix captures the spatial pattern of gamma-band co-activation in V1 and V4 that is due to common driving through stimuli at varying positions.

**Figure 4.**
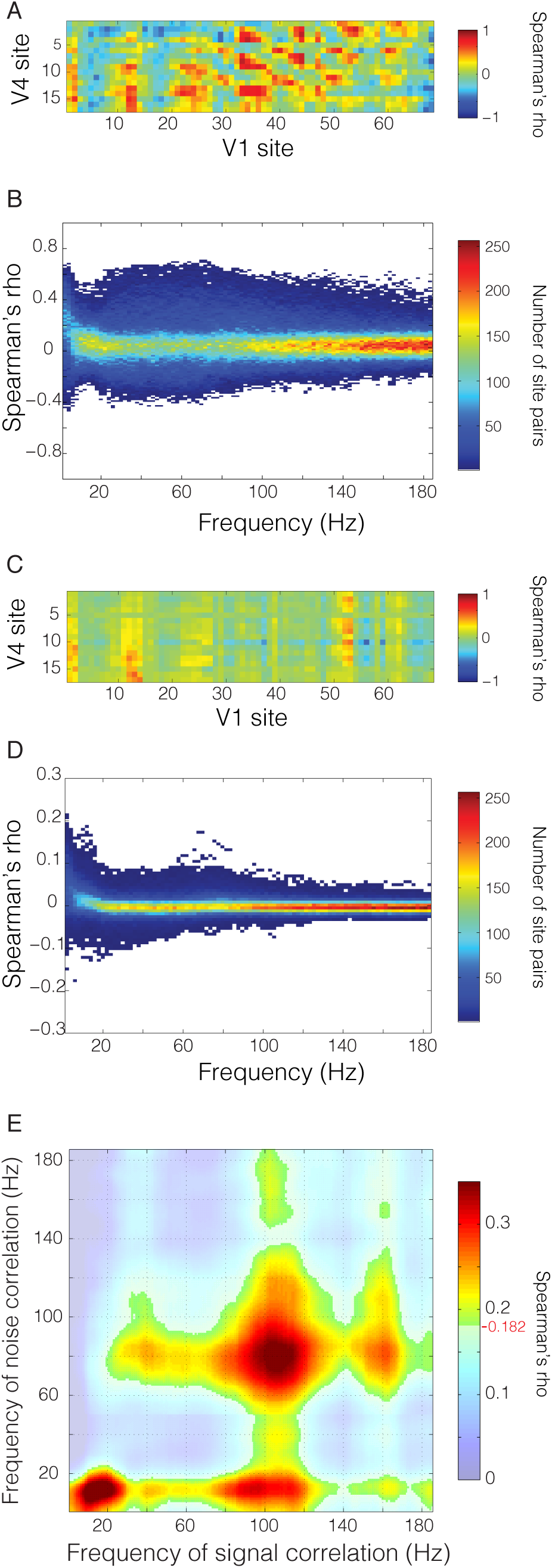
Signal and noise correlations. (A) Signal correlation matrix in the gamma band (80-95 Hz) for all V1-V4 site pairs. (B) Distribution of signal correlation values for all inter-areal site pairs as a function of frequency. (C) Noise correlation matrix in the same band as in (A). (D) Distribution of noise correlation values as a function of frequency. (E) Correlation of inter-areal signal and noise correlation matrices (panels A and C) across frequencies. Frequency-frequency plane threshold P<0.05 corrected for multiple comparisons. Significance threshold is denoted on colorbar by red line and value.

Figure 4B shows the distributions of signal correlation values across all V1-V4 site pairs as a function of the frequency for which the power was taken. While specific frequency bands contain most stimulus-related information, signal correlations occur across a broad spectrum of frequencies.

## Noise Correlations

As mentioned above, the difference between the average response and the actual response to a given stimulus presentation has been considered as noise. While we believe that this so-called noise component is actually not noise, but might reflect sources of uncontrolled endogenous variance. e.g. top-down influences, we will use the signal versus noise terminology for consistency with previous literature. This noise, i.e. the deviations from the average response, has often been found to be correlated across recording sites, i.e. there is noise correlation. We computed noise correlations in order to assess the degree of power covariation between visual regions that was independent of the stimulus and therefore intrinsically generated (sample site pair, same as above, shown in Fig. 3F). The noise correlation is complementary to the signal correlation because, for each pair of recording sites, it captures the intrinsic covariation in power across repeated trials of identical stimulation as opposed to shared stimulus selectivity. Noise correlation was computed for all V1-V4 site pairs and for each frequency (Fig. 3H). Figure 4C shows the matrix of noise correlation values from all possible V1-V4 site pairs for the gamma-frequency band (between 81 and 96 Hz) for monkey P. This matrix captures the spatial pattern of gamma-band co-activation in V1 and V4 that occurs during stimulation, but reflects intrinsic trial-by-trial variation.

Figure 4D shows the distributions of noise correlation values across all V1-V4 site pairs, and reveals similar characteristics as mentioned above for the signal correlations.

## Similarity of noise correlations to signal correlations

In order to test whether the spatial pattern of noise correlations resembled the spatial pattern of signal correlations, we calculated the correlation between those two metrics, across inter-areal site pairs. As mentioned above, since both, the signal and the noise correlations were determined as a function of frequency, and we considered all possible frequency pairs, this resulted in a frequency-by-frequency matrix of correlation values. We found that noise correlations correlated significantly with signal correlations in specific frequency ranges (Fig. 4E). These frequency ranges correspond to those showing the greatest stimulus selectivity (Fig. 1D). Thus, inter-areal power covariations that are extrinsically driven by different visual stimuli are mirrored in power covariations that occur intrinsically in the trial-by-trial variation around the stimulus response, and this effect is most prominent in the frequency ranges in which retinotopic selectivity is expressed.

The existence of frequency-specific similarity in the spatial pattern of inter-areal covariation cannot be trivially explained by the pattern of signal correlations. It is possible that noise correlations could occur in an unspecific manner, either spectrally or spatially. For example, on a given trial, all visual channels could have high or low power in a narrow or broad range of frequencies, in which case the degree of noise correlation could be the same as observed, but it would not reflect the underlying retinotopic organization. This would be the case if noise correlations reflected unspecific, global, signal variance. However, our results suggest that noise correlations between visual areas obey the functional organization of the underlying cortex.

## Similarity of spontaneous correlations to signal correlations

The signal and noise correlations investigated so far were derived from the same data, extracting signal-driven variance or endogenous variance and their respective correlation structure. We wondered whether the intrinsically generated covariance resembled the signal correlation also when it was taken from entirely independent data. To this end, we analyzed the fixation periods of separate recording sessions, mostly on different days than the recording sessions analyzed so far. During these fixation periods, the monkey sat quietly in the booth and fixated a central fixation point while awaiting a different task. We calculated correlations between spontaneous power fluctuations across time after concatenating trials, and we refer to them as spontaneous correlations (Fig. 3I and J). Figure 5A shows the spontaneous correlations across the complete matrix of V1-V4 site pairs for the gamma-frequency band in monkey P. Figure 5B shows the distributions of spontaneous correlation values across all V1-V4 site pairs as a function of the frequency for which the power was taken.

**Figure 5.**
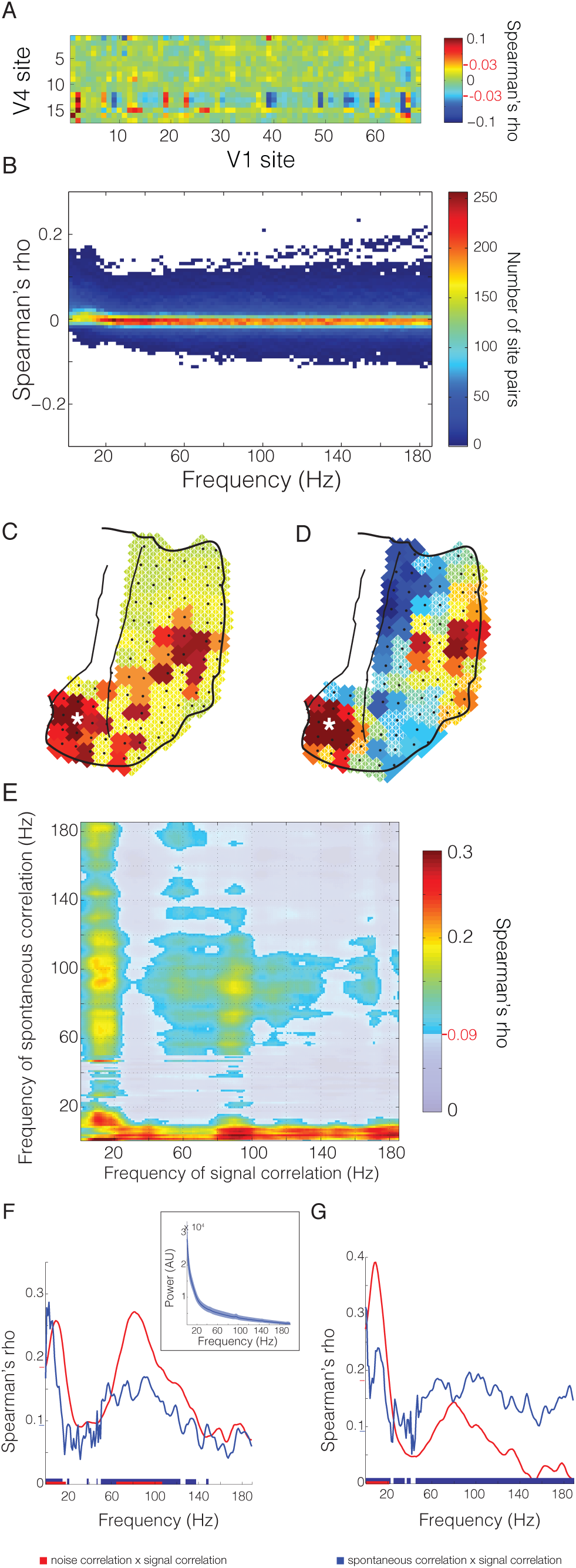
Spontaneous correlations. (A) Spontaneous correlation matrix in the gamma band (80-95 Hz) for all V1-V4 site pairs. (B) Distribution of spontaneous correlation values for all inter-areal site pairs as a function of frequency. Example topographies for spontaneous correlation from the marked V4 site across all visual channels in the (C) theta (6-8 Hz) and (D) gamma (80-95 Hz) bands (same color bar as (A)). Statistical significance is denoted by red lines and values on colorbar, as well as, white borders on non-significant values in topography. (E) Correlation of inter-areal spontaneous and signal correlation matrices (panel A from figures 4 and 5) across frequencies. Frequency-frequency plane threshold P<0.05 corrected for multiple comparisons. Significance threshold is denoted on colorbar by red line and value. (F) Comparison of intrinsic correlations with signal correlations for the gamma f requency band (80 - 100 Hz). The mean correspondences of spontaneous correlation (in blue) and noise correlation (in red) with signal correlation are shown to illustrate their similar spectral profiles. The line spectra are vertical cuts of the respective frequency-frequency plots shown in Figs. 4E and 5E). Significance threshold at P<0.05 corrected for multiple comparisons, illustrated in blue and red on the y-axis and significant bands of correspondence illustrated as blue and red bars along the x-axis. (inset) Spectrum of absolute power of the normalized raw signal (see Experimental procedures for details) for the activity during fixation used to compute spontaneous correlation. Mean across visual sites shown in dark blue. Shaded region denotes standard deviation across sites. (G) As in (F), but for the low frequency band (1-20 Hz). The mean correspondences of spontaneous correlation (in blue) and noise correlation (in red) with signal correlation are shown to illustrate their different spectral profiles. The line spectra are vertical cuts of the respective frequency-frequency plots shown in Figs. 4E and 5E). Significance threshold is as explained for (F).

Differences in the pattern of correlation for the low and high frequency activity were evident in the topographic distribution of spontaneous correlation values. Figure 5C illustrates the spatial pattern for the spontaneous correlation from a seed site in V4 in the 6-8 Hz band. The portion of V1 with the highest spontaneous correlation corresponded to the portion with the same retinotopic selectivity as the V4 seed site (cf. Fig. 2). This retinotopic correspondence held also for spontaneous power fluctuations in the gamma band (80-95 Hz, Fig. 5D). Whereas spontaneous correlation values for the theta band were almost exclusively positive, the values for the gamma band were both positive and negative, as expected from figure 5B.

We found spontaneous correlations to be correlated with signal correlations across V1-V4 site pairs. The retinotopically specific spontaneous coactivation showed a superposition of several correspondences (Fig. 5E), specifically: 1) between spontaneous and signal correlations that were both in either the gamma-band range, or the alpha/beta-band range 2) between signal correlations in the beta band and spontaneous correlations in a very broad band, 3) between spontaneous correlations in the theta/alpha band and signal correlations in a very broad band. The broadband correspondences suggest arrhythmic or 1/f processes as the underlying source. This superposition of rhythmic and arrhythmic correspondences was revealed through the investigation of the complete frequency-by-frequency plane. By investigating the correspondence between retinotopy and intrinsic signals at all frequency pairs, we were able to uncover important distinctions between passive (Fig. 5E) and active (Fig. 4E) states.

Signal correlations in the gamma band had a similar pattern of correspondence with spontaneous correlations as they had with noise correlations. In order to demonstrate this similarity, we plot the cross-sections showing the correlation spectra for spontaneous and noise correlation with the gamma-range signal correlation (i.e. Figs. 4E and 5E) in Fig. 5F. These cross-sections show clear peaks in the delta/theta band and the gamma band. The presence of such peaks for the spontaneous correlation, measured during passive fixation, is in stark contrast to the absolute power spectrum of the LFP during the fixation period, which shows a typical 1/f characteristic (Fig. 5F inset). Thus, although there were no appreciable spectral peaks, the spontaneous V1-V4 covariations revealed band-limited processes that were also specifically modulated in the two areas by localized stimuli. In contrast, signal correlations in the alpha/beta band had a different pattern of correspondence with spontaneous correlations versus noise correlations. This is demonstrated by corresponding cross-sections in Fig. 5G. Here, the broadband, arrhythmic component of the spontaneous correspondence diverges from the band-limited low frequency component, which is present both in the spontaneous correlations and the noise correlations.

## Spontaneous correlations recapitulate stimulus correlations

The retinotopic structure of inter-areal spontaneous correlations can be demonstrated directly in topographical form, as shown in Fig. 5C and D. In order to assess the degree of this correspondence, we investigated whether the inter-areal pattern of spontaneous correlations allowed us to derive an ordered topography from stimulation-free data, similar to the retinotopic map derived from visual stimulation data.

To this end, we grouped sites in V4 into regions of interest (ROI), when they shared similar selectivity either for eccentricity or for elevation. We calculated spontaneous correlations between each V4 ROI and all V1 sites, separately for the low (6-8 Hz) and gamma (80-95 Hz) frequency bands. We hypothesized that a given ROI, with selectivity for either a particular eccentricity or elevation would show strongest spontaneous correlation with V1 sites sharing the same stimulus selectivity. We indeed found this to be the case. We colored each V1 site according to the eccentricity (Fig. 6A and C show the maps for power in the 6-8 Hz and 80-95 Hz bands, respectively) or elevation (Fig. 6B and D, as in A and C) of the V4 ROI to which it showed the strongest spontaneous correlation. To quantify the similarity between these topographies and the retinotopic maps from figure 2, we computed the spatial correlation between them. We limited this spatial correlation analysis to sites in V1, because our selection of ROIs in V4 was already based on stimulus preference and so similarities inside V4 were trivial, whereas the pattern seen in V1 was solely the result of topographic specificity of the spontaneous correlations. Both maps derived from spontaneous activity were highly correlated with the retinotopic maps (6-8 Hz frequency band: Eccentricity map, Spearman’s rho= 0.366, False Discovery Rate across all frequencies, q<0.05; Elevation map, Spearman’s rho= 0.37, q 0.05. 80-95 Hz frequency band: Eccentricity map, Spearman’s rho = 0.274, q <0.05; Elevation map, Spearman’s rho = 0.266, q <0.05.). Further, when examining the spatial correlation of the maps derived from spontaneous activity with the retinotopic maps across all frequencies, we again found correspondence specifically in the band below 20 Hz and a broad gamma band between 60-150 Hz. These values correspond roughly to the correspondence found through noise correlations, spontaneous correlations, as well as the frequency bands showing the highest stimulus selectivity. Further, in the spectral region between the two zones of highest correspondence (20-30 Hz), we found some zones of negative correlation, suggesting a different pattern of inter-areal interaction occurring in this frequency range during spontaneous activity. This supports the previous results and further confirms that intrinsic power variations have spatial structure that recapitulates the functional topography of the underlying cortex.

**Figure 6.**
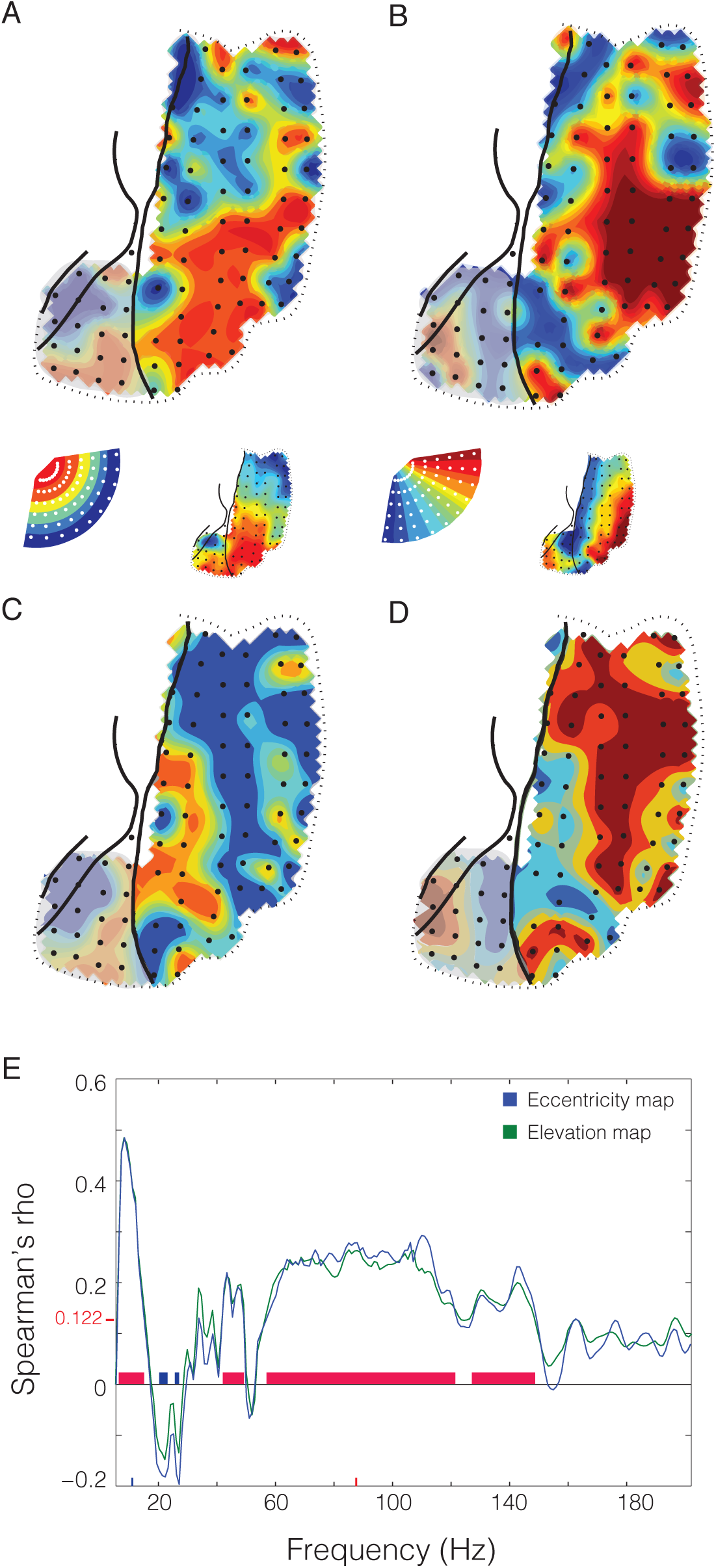
Retinotopic maps computed from the pattern of spontaneous correlation. (A and Map of eccentricity during passive fixation; each recording site is colored based on the mean eccentricity tuning of the two maximally correlated V4 sites during passive fixation in the (A) theta (6-8 Hz) and (C) gamma (80-95 Hz) band. (B and D) Map of elevation during passive fixation as in A and C. The retinotopic maps computed from gamma power as in Figure 2 are shown as insets for comparison. (E) Spectrum of spatial correlation values between spontaneous retinotopic maps and stimulus induced retinotopic maps in area V1. Blue line is for eccentricity, green for elevation. Significance threshold is denoted on y-axis by redline and value.

## Similarity of spontaneous coherence and directed influences with signal correlation

Given that both, noise correlations and spontaneous correlations, showed a similar spatial pattern as signal correlations, we asked whether this spatial pattern could also be found in the spontaneous inter-areal coherence, i.e. in a metric of phase synchronization during spontaneous activity recorded during pre-stimulus fixation periods. It is possible that the spectral power between areas could be correlated, but that the inter-areal signals would not exhibit a specific phase relationship. For example, rhythmic activity could be generated locally at two locations. The power, or other second-order characteristics, of such locally generated rhythms could be correlated, while no first-order dependencies, such as phase coherence, would be present between them. Such a case is particularly plausible in a scenario where the signal power does not depart from a 1/f spectrum, suggesting a lack of strong oscillatory activity. Despite this, inter-areal coherence spectra for the fixation period do demonstrate rhythmic components despite the lack of oscillatory structure in the power (Fig. S4). To our surprise, we found that inter-areal coherence between visual regions obeyed retinotopic organization in frequency bands similar to those showing similarity in noise correlations and spontaneous correlations (Fig. 7A, all spectra are compared in Fig. S5). Thus, even though no clear gamma peak was noticeable in the power spectrum, the correlation analysis revealed that there was spontaneous gamma-band coherence between V1 and V4 that occurred selectively between regions of retinotopic correspondence. As with the spontaneous correlations, derived from periods of passive fixation, the correspondence between spontaneous phase synchronization and retinotopic organization displayed broadband components in addition to band-limited components. These were mostly between retinotopic organization present in the activity below 20 Hz and spontaneously synchronized broadband activity.

**Figure 7.**
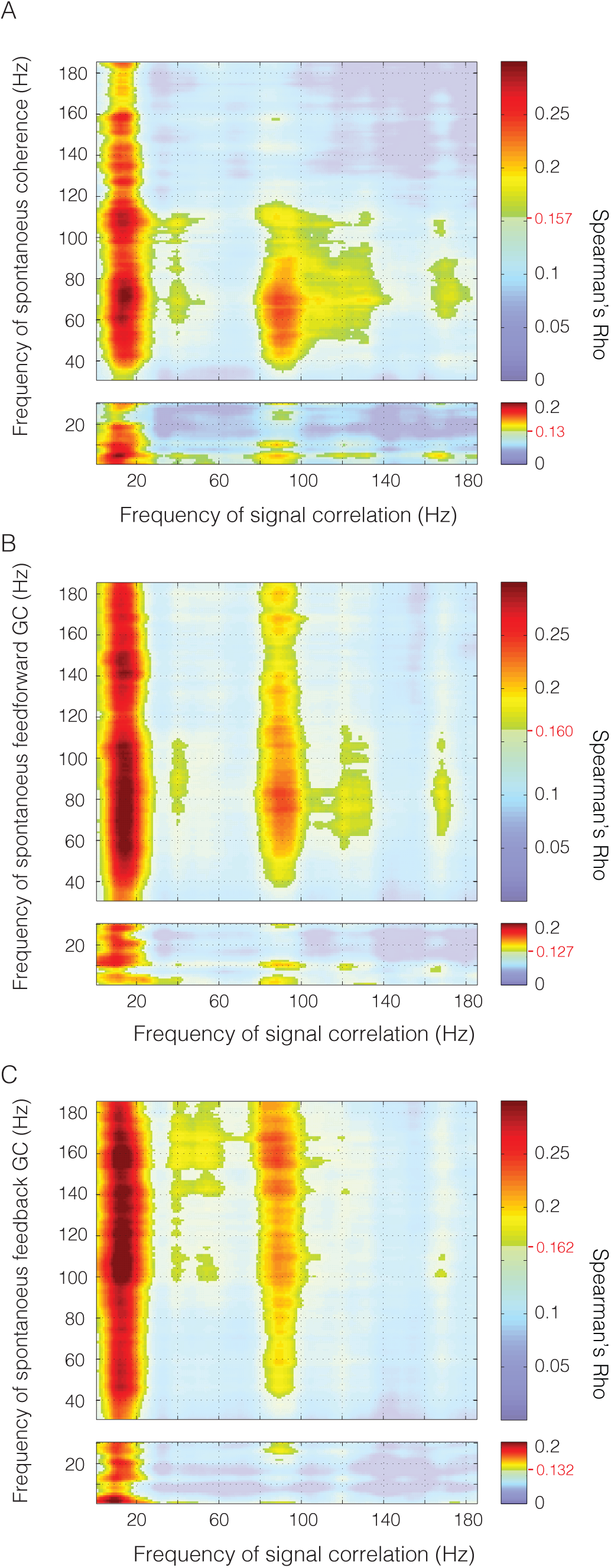
Spontaneous coherence and directed interaction. Correlation of (A) spontaneous coherence, (B) spontaneous feed-forward GC, and (C) spontaneous feed-back GC with signal correlation for all frequencies. Frequency-frequency plane threshold set at P<0.05 corrected for multiple comparisons. Significance threshold is denoted for each comparison by the red line and value on the colorbar.

Given that this was the case, we attempted to assess the directionality of the interaction by measuring the granger causal (GC) influences in both directions. We found that signal correlation determined the spatial structure of inter-areal GC influences both in the direction from V1 to V4 (Fig. 7B) and in the direction from V4 to V1 (Fig. 7C, inter-areal granger spectra are shown in Fig. S4).

## Discussion

In summary, we found intrinsic networks subserved by local and inter-areal synchronization, whose spatial structure correlates significantly with stimulus driven patterns of inter-areal co-activation. Concretely, we found that the spatial structure of signal correlations was recapitulated in 1.) the trial-to-trial covariability of stimulus induced power (noise correlations), 2.) the correlation of spontaneous power fluctuations during fixation (spontaneous correlations) and 3.) the spontaneous coherence and GC influences during fixation. We take the specificity of these intrinsically generated patterns of synchronization as a robust indicator of highly organized endogenous activity. Notably, the same spatial pattern of intrinsic covariation was present both during stimulation, i.e. in the noise correlation, and in the absence of stimulation, i.e. in the spontaneous correlation. The patterns present during passive, non-stimulated states were a superset of those present during stimulation (Luczak et al., 2009), suggesting that the intrinsic dynamics may differ between the two states. This suggests that in addition to the role intrinsic networks may play during passive states, the respective intrinsic networks may influence the pattern of activity during perception and action, and thereby, impact behavior. The temporal structure of the observed intrinsic networks showed both arrhythmic, broadband activity, as well as characteristic frequency bands, corresponding to the rhythms most specifically modulated by visual stimulation, namely the theta, alpha/beta and gamma rhythms.

One potential concern is that the signal correlation is correlated to intrinsic networks in a spectrally specific way only because the signal correlation itself is spectrally specific to begin with. Yet, a close examination of the data does not confirm this. For example, figure 4E shows peaks in the congruence between signal and noise correlation for signal correlations in the frequency bands of roughly 10-20 Hz and of 100-120 Hz. Figure 4B shows the distributions of signal correlations and reveals that signal correlations in the 10-20 Hz band are among the weakest, and signal correlations in the 100-120 Hz band are in the middle of a broad band of all frequencies above 40 Hz, for which the distribution of signal correlations gets progressively narrower. Thus, the frequency bands, in which the signal correlation is particularly congruent to intrinsic correlation patterns, are not particularly conspicuous in the spectral distribution of signal correlation values. Very similar arguments can be made for the congruence between signal correlation and spontaneous correlation, coherence or GC influences. The frequency bands of highest congruence do not correspond to frequency bands with particularly strong signal correlation. The same reasoning holds for the distributions of noise correlations and spontaneous correlations, which are shown in figures 4D and 5B, and also lack conspicuous structure in the ranges in which they show spatial congruency to the signal correlation. Furthermore, the statistical tests for establishing congruency between signal correlations and intrinsic networks rested on the inter-areal spatial correlation pattern. By randomizing the spatial relation between inter-areal sites, we generated surrogate distributions of congruency metrics. Importantly, this randomization left the underlying distribution of the signal correlation unchanged.

The frequency-specific interactions during spontaneous activity occurred in the absence of clear peaks in the LFP power spectrum. The retinotopically specific band-limited inter-areal correlation patterns were embedded within the 1/f frequency spectrum. This suggests that: the involved networks have the tendency to structure their activity in rhythmic modes even when they are not strongly activated, and 2) the conventional power spectrum is an insensitive tool to detect rhythmicity. This latter point is supported by other recent findings. For example, the relative phases between simultaneously recorded LFPs are highly structured in the gamma-frequency band, both during visual stimulation and in the pre-stimulus baseline, when there is no peak in gamma power (Maris et al., 2013). Furthermore, the spike times of putative inhibitory interneurons are selectively locked to the phase of activity in the gamma-band during the pre-stimulus baseline, again in the absence of any appreciable gamma power peak (Vinck et al., 2013).

Signal correlations showed spatial correspondence with noise correlations mainly for rhythmic components, while they showed correspondence with spontaneous correlations for both rhythmic and arrhythmic components. This difference suggests that cortical activity exhibits distinct dynamics during stimulation and in its absence. While the absence of stimulation might leave cortex to wander through a wider range of dynamical states, the presence of stimulation might constrain this range of dynamic states and thereby present with clear rhythms. In the absence of such stimulus-related constraints, local and short-lived rhythms might well be present, but with variable frequency, strength and spatial extension. Such passive states would therefore exhibit a characteristic 1/f spectrum (Freeman, 2004; He, 2014; Linkenkaer-Hansen et al., 2001). However, nested within an arrhythmic spectrum, highly specific patterns of organization, such as cross-frequency interactions (He et al., 2010) or topographically specific inter-areal coupling, may reveal specific rhythmic components. Such spontaneously occurring rhythms might couple between V1 and V4 in a retinotopically specific way most strongly, when they occur in particular frequency ranges. This could explain why we find broadband correspondences together with correspondences that peak in particular, well-known frequency bands.

Interestingly, the two bands of high correspondence between intrinsic covariation and stimulus driven covariation correspond well with previous findings. Activity in the gamma band is more selective for stimulus properties than oscillations in other bands (Frien and Eckhorn, 2000; Frien et al., 2000). Furthermore, when natural movies were shown to anesthetized monkeys, most information about the movies was contained in the power time courses of LFP components between 1-8 Hz and between 60-100 Hz (Belitski et al., 2008). These bands correspond roughly to the ones in which we find consistency between retinotopy and intrinsic covariation, suggesting that those frequency bands may generally be involved in the representation of visual features within areas and their communication between areas. In fact, within areas, stimulus representation is more accurate for the spikes that are optimally aligned to the gamma rhythm as opposed to spikes that occur at random phases of the gamma cycle (Womelsdorf et al., 2012). Similarly, gamma-band synchronization might facilitate the inter-areal communication of representations: 1) Inter-areal gamma phase locking leads to enhanced inter-areal interactions (Womelsdorf et al., 2007); 2) The selective inter-areal communication of attended stimuli is subserved by a corresponding selective inter-areal gamma-band synchronization (Bosman et al., 2012; Grothe et al., 2012). One interesting possibility is that the pattern of inter-areal coupling in the gamma frequency range is dependent on the phase of low frequency oscillations. Such phase-amplitude coupling has been demonstrated in many other data sets (Canolty, et al. 2006; He, et al., 2010) and could provide a link between the low and high frequency retinotopic correspondence we find in our data. These converging results indicate that the intrinsic patterns we report here may play a role in the brain’s endogenous sampling and routing of visual information.

Intrinsic networks defined by structured endogenous activity have been observed at many spatial and temporal scales. Luczak et al. investigated the similarity of stimulus driven and spontaneous patterns of population activity within auditory or somatosensory cortex of both anesthetized and awake rats (Luczak et al., 2009). They found that spike timing patterns were conserved across states of stimulation and spontaneous activity, and that the spatial patterns observed during stimulation constitute a subset of all spatial patterns visited by the network during spontaneous activity. Further, they demonstrated the similarity of pair-wise correlations between individual cell’s firing rates during stimulus driven and spontaneous activity as well as in noise correlations across repeated stimuli. Similar observations have been made for the entire map of activity across cat primary visual cortex (Arieli et al., 1995; Kenet et al., 2003; Tsodyks et al., 1999). The distribution of activity in the absence of visual stimulation resembled maps evoked by specific visual stimuli more frequently than expected by chance. Between monkey visual areas, retinotopically corresponding regions show spontaneous co-variations in the fMRI BOLD signal (Vincent et al., 2007). Similar inter-areal intrinsic networks of BOLD covariation have been found abundantly in human subjects and have been related to different functional brain networks. The finding of intrinsic, whole brain networks with fMRI has expanded the scope of patterned spontaneous activity from the level of sensory maps to general inter-regional interactions. Further, the ability to investigate intrinsic networks in humans has led to studies that directly demonstrate the functional role of these networks: Human intrinsic networks have been shown to be shaped by experience and to affect neuronal and behavioral responses. Related effects of experience have also been shown for local circuits in animals (Berkes et al., 2011). These results expand the implications of intrinsic networks and illustrate their relevance for understanding both healthy and diseased brain function.

Electrophysiological correlates of intrinsic networks observed with BOLD have been sought in order to begin bridging the gap between detailed accounts of endogenous dynamics within cortical areas to the patterns of activity observed with fMRI across the whole brain. In visual cortex, BOLD signal fluctuations show a positive correlation primarily with LFP power in the gamma frequency range (Logothetis et al., 2001; Mukamel et al., 2005; Niessing et al., 2005; Scheeringa et al., 2011; Schölvinck et al., 2010). LFP power in the gamma frequency range can reflect both, rhythmic neuronal synchronization at the gamma rhythm (Eckhorn et al., 1988; Fries et al., 2001; Gray et al., 1989) and the broadband spectral signature of basic biophysical processes like spikes and/or postsynaptic potentials (Einevoll et al., 2013; Miller et al., 2009), a distinction that is increasingly made explicit (Ray and Maunsell, 2011). The BOLD signal appears related to both, the strength of broadband gamma-range power (Mukamel et al., 2005) and the strength of the band-limited gamma rhythm (Scheeringa et al., 2011; Schölvinck et al., 2010). Correspondingly, LFP power in the gamma-frequency range is correlated between corresponding regions of the two hemispheres (Nir et al., 2008), fluctuates spontaneously according to tonotopic maps in auditory cortex (Fukushima et al., 2012), and reflects regional boundaries in somatomotor cortex, in close correspondence to BOLD signal fluctuations (He et al., 2008). When activity in BOLD-signal defined intrinsic networks is directly correlated to EEG band-limited power, distinct networks relate to various frequency bands (Mantini et al., 2007). Conversely, power fluctuations in different frequency bands of the EEG signal are related to distinct spatial patterns of brain activation shown with simultaneous fMRI (Laufs et al., 2003a; 2003b). Power co-fluctuations were also demonstrated directly with source-projected MEG, revealing distinct spatial networks for power in different frequency bands (de Pasquale et al., 2010; Hawellek et al., 2013; Hipp et al., 2012). Related findings have been reported in recordings from cat brain regions homologous with two well-described human fMRI intrinsic networks, the task-on and task-off networks (Popa et al., 2009). Between those two networks, total LFP power during spontaneous activity showed an overall difference, and periods in which band-limited power time courses were anti-correlated. In the task-off network, reduced power was surprisingly correlated to enhanced neuronal firing rates, illustrating the importance of combined multi-level investigations.

We have shown that the intrinsic networks are subserved by specific patterns of synchronization. The pattern of synchronization varies between dominantly narrow-band, rhythmic activity during periods of stimulation and a mixture of both broad- and narrow-band synchronization during periods of spontaneous activity. Rhythmic activity was evident even when the power spectrum exhibited no spectral structure. Rhythmic and arrhythmic activity occurred in distinct portions of the frequency-frequency interactions, suggesting that they occur distinctly within the data. Rhythmic activity was evidenced by clear spectral peaks that exclude broadband power changes as underlying causes and rather correspond to classical frequency bands. Arrhythmic activity was mainly present in the correspondence of intrinsic signals with retinotopic organization below 20 Hz. The segregation of these two modes of activation suggest that broadband activity may be specifically modulated by low frequency activity, as has been demonstrated in human ECoG recordings (Canolty et al., 2006; He et al., 2010). The rhythmic activation of particular interest occurred in the gamma-frequency band and has been linked to inter-areal communication. Communication is likely supported by gamma-band coherence. Surprisingly, gamma-band coherence, normally considered to exist exclusively during stimulus driving, was found in spontaneous activity to reflect the spatial pattern of signal correlations. This finding provides a potential link from the putative mechanism of communication-through-coherence to the organization of endogenous activity in intrinsic networks. The link could be shown due to ECoG recordings that combined the high temporal and high spatial resolution with coverage of two visual areas. The high temporal resolution was necessary to reveal the spectral specificities, the high spatial resolution allowed to reveal the topographic specificity and the coverage was required to allow the inter-areal correlation between spatial patterns of spontaneous and stimulus driven activity. It will be an intriguing possibility for future research to capitalize on these features and investigate whether e.g. the influence of intrinsic networks on stimulus responses or behavior is spectrally specific, or whether experience shapes intrinsic networks particularly or differentially in different frequency bands. Furthermore, it will be crucial to extend coverage to both hemispheres to allow investigation of bilateral symmetric patterns, which have been the hallmark of many fMRI demonstrations of intrinsic networks, and finally to add single cell recordings that will allow the integration of population dynamics into large-scale patterns.

## Experimental Procedures

### Neurophysiological Recording Techniques and Signal Preprocessing

All procedures were approved by the ethics committee of the Radboud University, Nijmegen, Netherlands. Neuronal recordings were made from two left hemispheres in two monkeys through a micro-machined 252-channel electrocorticogram-electrode array with 1mm diameter contacts spaced by 2.5-3mm and implanted subdurally (Bosman et al., 2012; Brunet et al., 2013; 2014; Rubehn et al., 2009; 2014). Electrode impedance was ∼3 kΩ at 1kHz. The reference electrode was a silver ball places over the contralateral visual cortex. The common recording reference was removed through calculating the bipolar derivation before any further processing. Briefly, a 6.5 × 3.4 cm craniotomy over the left hemisphere in each monkey was performed under aseptic conditions with isoflurane/fentanyl anesthesia. The dura was opened and the ECoG was placed directly onto the brain under visual control. Several high-resolution photos were taken before and after placement of the ECoG for later co-registration of ECoG signals with brain regions. After ECoG implantation, both the bone and the dura flap were placed back and secured in place. ECoG electrodes covered numerous brain areas, including parts of areas V1, V2, V4 and TEO. As mentioned in the main text, retinotopic mapping revealed two contiguous maps of space, one behind the lunate sulcus for areas V1/V2, and another one between the lunate and the superior temporal sulcus for areas V4/TEO. For simplicity, we will refer to ECoG sites in the V1/V2 map as V1, and to sites in the V4/TEO map as V4. After a recovery period of approximately 3 weeks, we started neuronal recordings. Signals obtained from the electrode grid were amplified 20 times by eight Plexon headstage amplifiers, then low-pass filtered at 8 kHz and digitized at 32 kHz by a Neuralynx Digital Lynx system. LFP signals were obtained by low-pass filtering at 200 Hz and down-sampling to 1 kHz. Power line artifacts were removed by digital notch filtering. The actual spectral data analysis included spectral smoothing that rendered the original notch invisible.

### Visual Stimulation

Stimuli and behavior were controlled by the software CORTEX. Stimuli were presented on a cathode ray tube monitor at 120 Hz non-interlaced. When the monkey touched a bar, a gray fixation point appeared at the center of the screen. When the monkey brought its gaze into a fixation window around the fixation point (0.85 radius in monkey K; 1 radius in monkey P), a pre-stimulus baseline of 0.8 s started. If the monkey’s gaze left the fixation window at any time, the trial was terminated.

Several sessions (either separate or after attention-task sessions) were devoted to the mapping of receptive fields, using 60 patches of drifting grating, as illustrated in Figure 1B. Gratings were circular black and white sine waves, with a spatial frequency of 3 cycles/degree and a speed of 0.4 degrees/s. Stimulus diameter was scaled between 1.2 and 1.86 degrees to account for cortical magnification factor. Receptive field positions were stable across recording sessions (see Figure S1D of Bosman et al., 2012).

### Data Analysis General

All analyses were performed in MATLAB (MathWorks) using FieldTrip (Oostenveld et al., 2011) (http://fieldtrip.fcdonders.nl). We calculated local bipolar derivatives, i.e., differences (sample-by-sample in the time domain) between LFPs from immediately neighboring electrodes. We refer to the bipolar derivatives as “sites.” Bipolar derivation removes the common recording reference, which is important when analyzing power correlations and/or coherence. In this scheme, neighboring sites have an electrode in common. We limited our analysis to inter-areal site pairs, which never have a common electrode. Subsequently, per site and individual epoch, the mean was subtracted, and then, per site and session, the signal was normalized by its standard deviation. These normalized signals were pooled across sessions with identical stimulus and task, unless indicated otherwise. In order to select solely visually selective recording sites, an ANOVA was computed across frequencies (Fig. 1C, Fig. S1B).

Time series of band-limited power were computed for each trial by filtering the data with a one pass, causal Butterworth filter of order 2. We used a one-pass filter in order to avoid stimulation-related activity from contaminating epochs of passive fixation. Qualitatively similar results were found by using shorter time periods and a two-pass filter. Data were filtered into 185 frequency bands with center frequencies between 2 Hz and 200 Hz. We used different pass bands in computing the analysis to confirm the robustness of the results, and in the results presented here, we chose three different pass bands for three different groups of center frequencies (cf) : 3 Hz for low (cf = 2-11 Hz), 6 Hz for medium (cf = 13-53 Hz) and 15 Hz for high (cf = 59-200 Hz). After filtering, we calculated the Hilbert transform and took the absolute value of the analytic signal. All fixation trials were then concatenated into a pseudo-continuous time series for calculation of the inter-areal correlation coefficient.

### Retinotopic maps

In order to construct retinotopic maps for eccentricity and elevation, the mean power was computed across trials for each stimulus location. First, each trial was cut to the time between 0.3 s after stimulus onset (to avoid the early transient activity) and 0.9 s. Frequency analysis was performed by computing the fast Fourier transform of the entire stimulation epoch. Each site was assigned an eccentricity and an elevation based on the position of the stimulus giving the maximal response in the gamma band. The overall pattern of retinotopic organization was robust to different manners of computing the preferred eccentricity and elevation and so the maximal stimulus was chosen because of its simplicity.

### Signal and Noise Correlations

In order to compare the spatial patterns of stimulus-driven correlations and stimulus-independent correlations, frequency-resolved signal and noise correlations were computed. Signal correlation was computed by first calculating the mean spectral responses per stimulus position, across trials, and then determining the Spearman rank correlation across the mean stimulus responses, between inter-areal site pairs. Noise correlation was computed by first calculating the trial-by-trial deviation from the mean response for each condition, and then determining the Spearman rank correlation across trials, between inter-areal site pairs. Noise correlations were computed independently for each stimulus position and then averaged across conditions in order to avoid any modulation by stimulus preference from contaminating the results. Across all inter-areal site pairs, we then calculated the Spearman rank correlation between signal correlations and noise correlations. The Spearman rank correlation was used to avoid assumptions about underlying distributions. Yet, results were essentially the same when Pearson correlation coefficients were used.

### Spontaneous correlation

For the analysis of spontaneous activity, we used the period of passive fixation during an attention task, which contained stimuli at two fixed positions, one of them contralateral to the recorded left hemisphere. Recordings during the attention task occurred in different sessions and often on different days than retinotopic mapping. We defined fixation as the time period from 0.3 s after fixation point onset and after the monkey acquired fixation until 0.1 s before the first stimulus appeared on the screen. As with signal and noise correlations, spontaneous correlations during fixation where computed for each frequency of interest and for each channel pair.

### Spontaneous Retinotopy

Retinotopic maps were computed during passive fixation by first determining the eccentricity and elevation preference of recording sites in V4. Once this was determined, V4 sites were grouped into regions of interest (ROIs) with similar eccentricity (6 ROIs) or elevation preference (10 ROIs). For each ROI, average time series of band-limited power were computed and correlated separately with power time courses of each V1 site. This resulted in 6 V1 maps of correlation for eccentricity and 10 maps of correlation for elevation. In order to compute the topography in V1 during the period of spontaneous activity, each recording site in V1 was assigned the value of the retinotopic preference of the V4 ROI it was most strongly correlated with. This resulted in maps of intrinsically generated retinotopy for each frequency of interest. In order to quantify the extent to which spontaneous retinotopy corresponded with stimulus driven retinotopy in area V1, we computed the spatial correlation of those maps computed during spontaneous activity with stimulus derived retinotopic maps for eccentricity and elevation.

### Spontaneous Coherence and Directed Influence

For the analysis of spontaneous coherence and directed influence, we used the same passive fixation periods as during spontaneous correlation. We computed the coherence and Granger causality (GC) using two separate analysis approaches, one for frequencies less than or equal to 30 Hz and the other for frequencies between 31 Hz and 190 Hz. We did this because high frequency components characteristically have less power and broader spectral bands than low frequency components. Therefore, it is advantageous to apply different frequency-domain smoothing to these two distinct frequency bands. For both analyses, we cut the fixation data into 500 ms segments and computed the cross-spectral density matrix (CSD) using the fast Fourier transform. In the low frequency band, we used a Hann taper. For the high-frequency band, we used multi-tapering with 11 tapers from the discrete prolate spheroid sequence (Mitra and Pesaran, 1999). This gave us a spectral smoothing of +/- 11 Hz. After calculation of the CSD, we either calculated coherence or GC. GC was calculated using non-parametric spectral factorization (Dhamala et al., 2008). The pattern of inter-areal coherence and GC was then correlated with the pattern of stimulus selectivity for each combination of frequencies.

### Statistical Testing

Wherever possible, data from both monkeys were combined. This amounts to a fixed-effect analysis for our sample of two animals. The false positive rate was controlled by our statistical testing procedure as follows. In order to compute significance thresholds for both, frequency planes and line spectra, surrogate data were used which broke the spatial pattern of inter-areal interaction but kept all other features (e.g. the distribution and inter-areal pattern of signal correlations and the distribution of intrinsic correlation and coherence) unchanged. This was done by creating 1000 random pairings of inter-areal site pairs and then computing the correlation across site pairs, between the signal correlations and 1) the noise correlation, 2) the spontaneous correlation, 3) the coherence, and 4) the GC influences. Each realization of the randomized structure resulted in a distribution of correlation coefficients. In all cases we set our threshold for significance across all tests at P<0.05. In the case of noise correlation, since this came from the same dataset as the index for stimulus selectivity, we used an omnibus based multiple comparison correction (Maris et al., 2007; Nichols and Holmes, 2002). In this case, we retained the maximum surrogate correlation coefficient (absolute value) across all combinations of frequencies and corrected our empirical distributions by thresholding at the P<0.05 level from this distribution of surrogate maxima. In the other cases, since the analysis were from independent data, we used the less conservative False Discovery Rate correction (Genovese et al., 2002). To compute FDR corrections, we retained all correlation values across all randomizations in a single distribution and took our significance threshold from the P<0.05 level of this complete distribution.

## ACKNOWLEGMENTS

We thank M. Corbetta for discussions during the genesis of this project, and R. Oostenveld for assistance in technical matters.

This work was supported by Human Frontier Science Program Organization grant RGP0070/2003 (P.F.), The Volkswagen Foundation Grant I/79876 (P.F.), the European Science Foundation European Young Investigator Award Program (P.F.), the European Union (HEALTH-F2-2008-200728, Brain-Synch to P.F. and C.L.), the LOEWE program (“Neuronale Koordination Forschungsschwerpunkt Frankfurt” to P.F.), the Smart Mix Programme of the Netherlands Ministry of Economic Affairs and the Netherlands Ministry of Education, Culture and Science (BrainGain to P.F. and C.L.), the European Commission Seventh Framework Programme (FP7/2007-2013) under grant agreement 604102 (Human Brain Project to P.F. and C.L.), and The Netherlands Organization for Scientific Research grants 452-03-344 (P.F.) and 016-071-079 (T.W.).

### COMPETING FINANCIAL INTERESTS

The authors declare no competing financial interests.

